# Learning Optimal White Matter Tract Representations from Tractography using a Deep Generative Model for Population Analyses

**DOI:** 10.1101/2022.07.31.502227

**Authors:** Yixue Feng, Bramsh Q. Chandio, Tamoghna Chattopadhyay, Sophia I. Thomopoulos, Conor Owens-Walton, Neda Jahanshad, Eleftherios Garyfallidis, Paul M. Thompson

**Affiliations:** Imaging Genetics Center, Keck School of Medicine, University of Southern California, Marina del Rey, CA, United States; Department of Intelligent Systems Engineering, Indiana University Bloomington, Bloomington, IN, United States

**Keywords:** diffusion MRI, tractography, variational autoencoder, anomaly detection, Alzheimer’s disease

## Abstract

Whole brain tractography is commonly used to study the brain’s white matter fiber pathways, but the large number of streamlines generated - up to one million per brain - can be challenging for large-scale population studies. We propose a robust dimensionality reduction framework for tractography, using a Convolutional Variational Autoencoder (ConvVAE) to learn low-dimensional embeddings from white matter bundles. The resulting embeddings can be used to facilitate downstream tasks such as outlier and abnormality detection, and mapping of disease effects on white matter tracts in individuals or groups. We design experiments to evaluate how well embeddings of different dimensions preserve distances from the original high-dimensional dataset, using distance correlation methods. We find that streamline distances and inter-bundle distances are well preserved in the latent space, with a 6-dimensional optimal embedding space. The generative ConvVAE model allows fast inference on new data, and the smooth latent space enables meaningful decodings that can be used for downstream tasks. We demonstrate the use of a ConvVAE model trained on control subjects’ data to detect structural anomalies in white matter tracts in patients with Alzheimer’s disease (AD). Using ConvVAEs to facilitate population analyses, we identified 6 tracts with statistically significant differences between AD and controls after controlling for age and sex effect, visualizing specific locations along the tracts with high anomalies despite large inter-subject variations in fiber bundle geometry.

## 1. INTRODUCTION

Whole-brain tractography based on diffusion MRI is commonly used to study white matter pathways in a variety of neurological and psychiatric conditions, including Alzheimer’s disease and Parkinson’s disease.^1–3^ Each whole-brain tractogram can generate between 500,000 and more than one million streamlines, making it computationally expensive to perform downstream analyses, such as tractogram filtering,^4^ bundle labeling,^5, 6^ and population analyses.^3^ Data reduction, segmentation and labeling of whole brain tractograms are valuable to accelerate large-scale studies of brain disease. Deep learning methods in particular may offer new ways to efficiently represent the streamlines and their normal range of variations, as well as detect anomalies in individuals and patient groups.

*Representation learning* uses machine learning, and deep neural networks in particular, to distill information from large datasets with rich features into a lower dimensional latent space. These lower-dimensional models are often constructed to satisfy specific objectives, such as principal components analysis (PCA), which creates an orthogonal linear matrix, or sequence of linear projections that accounts for the maximum amount of variance in the data,^7^ or autoencoders (AE), which compress and encode the original data using nonlinear mappings for more convenient visualization, sparse reconstruction, and even denoising. After training deep networks to create these mappings, the low dimensional representations, or embeddings, can be directly used or fine-tuned for downstream prediction tasks, such as disease classification or prognosis^8^ or domain-specific tasks such as outlier detection in tractography.^9^ Zhong *et al*., for example, encoded streamlines with a recurrent autoencoder and used the embeddings for bundle parcellation.^10^ A similar approach, using a convolutional autoencoder, was developed for tractogram filtering.^4^ While these studies show that the learned embeddings can retain bundle information such as their shapes and positions, the latent space for standard autoencoders is not continuous and the model is often prone to overfitting,^11^ making it difficult to evaluate embeddings on unseen data, and use them for population analyses which involve large amounts of data. Variational autoencoders (VAEs), on the other hand, are generative models that learn a mapping to a space of continuous latent variables, enabling sampling and interpolation, and they are more stable in practice. The latent space carries relevant information about the input data,^8^ thus model design, hyperparameter tuning, validation task design and interpretation of the latent space should be informed by domain knowledge.

In this study, we learn optimal low-dimensional embeddings of white matter tracts with a Convolutional VAE (ConvVAE),^12^ and apply this more compact representation to detect detect disease effects in tract geometry. Meaningful embeddings should ideally preserve distance metrics from the original streamline space, as well as inter-bundle distances; there is a vast literature on embeddings that are approximately distance preserving, including multidimensional scaling and distance-preserving autoencoders, and embeddings that also satisfy related conditions involving the distributions of distances between point pairs (e.g., *t*-SNE, UMAP, and more complex methods based on persistent homology and computational topology). We investigate how the dimension of the latent space affects the quality of the embeddings using distance correlation analysis. We further demonstrate the use of our ConvVAE for anomaly detection in group analysis of Alzheimer’s disease (AD) at the tract level. The benefit of creating geometrically consistent latent and input data spaces is that anomaly detection and discriminative tasks can then be tackled in the much lower-dimensional latent space.

## 2. METHODS

### 2.1 Data

We computed whole-brain tractography from 3D multi-shell diffusion MRI scans of 141 participants in the Alzheimer’s Disease Neuroimaging Initiative (ADNI)^13^ (age: 55-91 years, 80F, 61M) scanned on 3T Siemens scanners. The dMRI data consisted of 127 volumes per subject; 13 non-diffusion-weighted *b_0_* volumes, 6 *b*=500, 48 *b*=1,000 and 60 *b*=2,000 s/mm^2^ volumes with a voxel size of 2.0×2.0×2.0 mm. Participants included 10 with dementia (AD), 44 with mild cognitive impairment (MCI), and 87 cognitively normal controls (CN). dMRI volumes were pre-processed using the ADNI3 dMRI protocol, correcting for artifacts including noise,^7, 14–16^ Gibbs ringing,^17^ eddy currents, bias field inhomogeneity, and echo-planar imaging distortions.^18–20^ We applied multi-shell multi-tissue constrained spherical deconvolution (MSMT-CSD)^21^ and a probabilistic particle filtering tracking algorithm^22^ to generate whole-brain tractograms. Thirty white matter tracts were extracted from all subjects in the MNI space using DiPy’s^14^ auto-calibrated RecoBundles.^3, 6^

### 2.2 Model

Variational Autoencoders (VAEs)^23^ retain the basic structure of autoencoders: an encoder, decoder and latent space serving as an information bottleneck. The ConvVAE model encoder has 3 convolutional blocks, each comprised of 1D convolutional, ReLU activation, batch normalization and average pooling layers.^4^ The decoder mirrors the encoder architecture with deconvolution instead of convolution, and upsampling instead of pooling layers (see Figure 1). Since the convolutional layers are designed to accept only fixed-dimension inputs, streamlines modeled as a sequence of 3D points are either downsampled or upsampled to generate 255 equal length segments, connecting the point sequence, *s* = {*p*_1_, *p*_2_,…*P*_256_}. The ConvVAE was trained on bundles from 10 control subjects with a batch size of 512 streamlines using the Evidence Lower Bound (ELBO) loss, consisting of a reconstruction and regularization term to enforce constraints on the latent space. We used the Adam optimizer^24^ with a learning rate of 0.0002 and weight decay of 0.001, and trained the model for 100 epochs. Gradient clipping^25^ by L^2^ norm was applied to prevent vanishing gradients, with a max norm value of 2.

**Figure 1.**
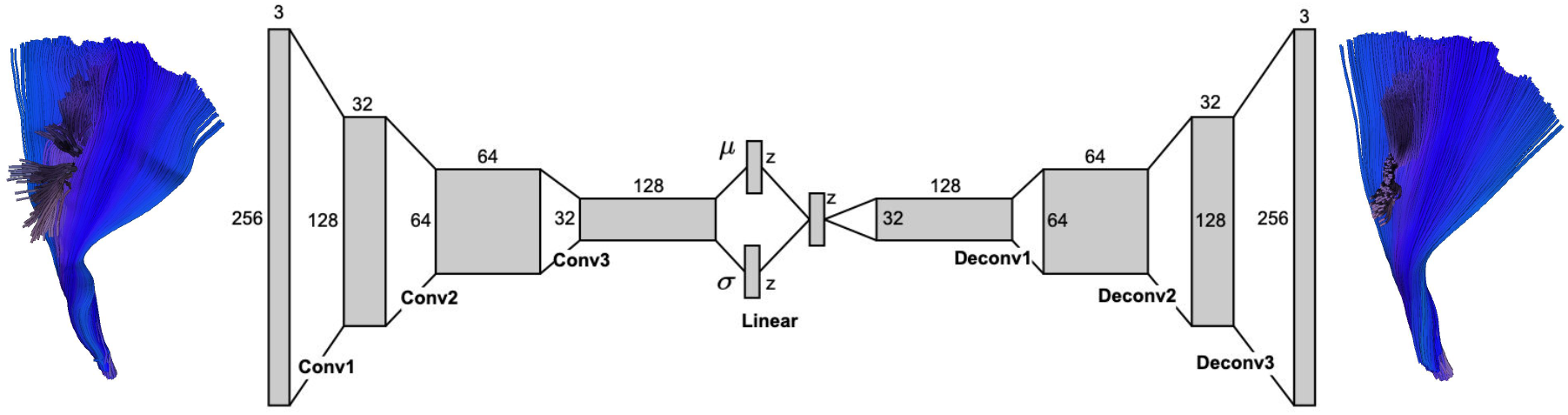
ConvVAE model architecture. The output shapes for one input streamline are indicated for each convolution and deconvolution block, consisting of 1D convolution, ReLU activation, batch normalization and average pooling layers. The embedding dimension *z* is a hyperparameter tuned to optimize the distance correlations between pairs of elements in the original and embedding space, as specified in Section 2.3.

### 2.3 Distance Preservation

To evaluate the quality of embeddings learned from ConvVAE, we conducted distance correlation analysis to see how well distances in the low dimensional latent space translate to distances in the observed streamline space. Multiple ConvVAE models were trained for 9 embedding dimensions *N_z_*, ranging from 2 to 32. We randomly sampled 300 streamlines from the training set, and computed their pairwise Euclidean distances between embeddings, as well as pairwise minimum direct flip (MDF) distance^26^ between the input streamlines. The correlation between these two distance metrics was evaluated using the Spearman’s rank correlation coefficient, Pearson correlation coefficient, and the coefficient of determination, *R*^2^, from linear regression. Since we expect that a zero distance in the latent space should correspond to zero distance in the streamline space, linear regression was fitted without an intercept. Repeating this correlation analysis for nine *N_z_* values, we plotted all three metrics to pick the maximum or elbow points 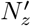 as the best latent dimension and used its corresponding model in subsequent tasks.

To further understand how well ConvVAE preserves *global* structure - which is a significant challenge for other dimension reduction methods such as *t*-SNE and UMAP - we conducted similar distance correlation analysis at the bundle level, using the ConvVAE model with embedding dimension of 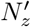 for a randomly selected training subject. Streamlines in bundle *b* are denoted by *X*(*b*) for *b* = 1, 2,…30, and embeddings corresponding to each bundle are denoted *Z*(*b*) for b =1, 2,…30. We used the same metrics to calculate pairwise inter-bundle centroid distances - MDF distance for streamlines bundle centroids Centroid(*X*(*b*)) calculated using QuickBundles,^26^ and Euclidean distance for embedding centroids Centroid(*Z*(*b*)) corresponding to bundle labels. We extend single centroid streamline analysis to bundles, using Bundle-based Minimum Distance (BMD)^27^ for inter-bundle distances between *X*(*b*), and Wasserstein distance^28^ for inter-bundle embedding distances between *Z*(*b*). The Mantel test^29^ was used to compute correlation between pairwise distance matrices, see Table 1.

**Table 1.** Results for bundle distance correlation analysis between bundles in the streamline space and embeddings in the latent space.

| Streamline Space |  | Latent Space |  | Mantel Test |  |
| --- | --- | --- | --- | --- | --- |
| Data | Distance Metric | Data | Distance Metric | Correlation $r_m$ | $p$-value |
| Centroid($X(b)$) | MDF Distance | Centroid($Z(b)$) | Euclidean Distance | 0.98 | 0.01 |
| $X(b)$ | BMD Distance | $Z(b)$ | Wasserstein Distance | 0.84 | 0.01 |

### 2.4 Anomaly Detection

Autoencoder-based approaches have been used in unsupervised anomaly detection for medical images.^31, 32^ The generative nature of the ConvVAE makes it possible to perform inference on new data, and any point in the smooth latent space can generate meaningful decodings instead of only minimizing reconstruction loss. In the context of anomaly detection when incoming data can have high variability and project to points on the latent space far from those of the training data, this quality of ConvVAE allows us to use the decoded output for outlier rejection and denoising. For this reason, ConvVAE may be less sensitive to outliers when the tractography generated is less than ideal.

To demonstrate the use of our ConvVAE model in group analysis for AD, we conducted an anomaly detection analysis between control, MCI and AD subjects at the bundle level. Since the ConvVAE was trained on bundle data from healthy control subjects, we expect its latent space to encode their relevant structural features, and the discrepancy between the reconstruction and the input (i.e., the reconstruction error) when the model is applied to new subjects can be used as a metric to flag anomalies. Chamberland *et al*.^33^ used autoencoders trained on dMRI microstructural features, and derived anomaly scores for individuals using the mean absolute error (MAE) between the input and reconstructed features.

Here we calculate MAE scores per bundle for all subjects excluding those used for training (CN:77, MCI:44, AD:10) and control for age and sex effect using linear regression.^33^ We stress that in this application, we do not aim to detect microstructural differences related to disease, but deviations in fiber tract geometry, and geometric distortions that may arise due to brain atrophy. Group MAE scores are calculated using a weighted average from individual MAE scores. Two-tailed independent sample t-tests assuming equal variance were performed at *α* = 0.05 between control and MCI (CN-MCI), and control and AD (CN-AD) groups. The Benjamini-Hochberg false discovery rate (FDR) correction was applied to adjust for multiple comparisons for all tracts. To further understand the group structural differences along each bundle, we calculated MAE of 100 segments along the length of the bundles per subject. The segments or assignment maps along the length of the bundles are computed using BUAN.^3^ We then regress the mean MAE across all segments on age and sex using linear regression. For the tracts that were significant in either of the previous CN-MCI and AD-MCI comparisons, two-tailed t-tests with FDR correction were also performed at each segment *s*_1_, *s*_2_,…*s*_100_ along the tract.

## 3. RESULTS

### 3.1 Embeddings Evaluation

To evaluate the effect of embedding dimensions on the latent space, we plot distance correlation from 300 subsamples with Euclidean embedding distances on the x-axis and streamline MDF distances on the y-axis, for embeddings of dimension *N_z_* = 2, 6, 8, 16, see Figure 2. A linear fit with zero intercept is plotted in red dashed line, and the Spearman *r*, Pearson *r* and *R*^2^ are indicated in the legend. We can see from plot N_z_ = 2 that the distance relationship doesn’t follow a linear relationship as reflected by the poor R^2^ score, where streamline distances correspond to a smaller range of embedding distances than those at higher *N_z_*. As *N_z_* increases, embedding distances are more strongly correlated with streamline distances and more closely follow a linear relationship. Unlike *N_z_* = 6 and 8 however, the *N_z_* = 16 plot shows that zero streamline distances correspond to non-zero embedding distances, resulting in low *R^2^* score. In Figure 3, we plot all three metrics against *N_z_* for all 9 models, and found that the ConvVAE model with *N_z_* = 6 has the best distance correlation and *R^2^*, indicating that streamline distances are best preserved at this dimension.

**Figure 2.**
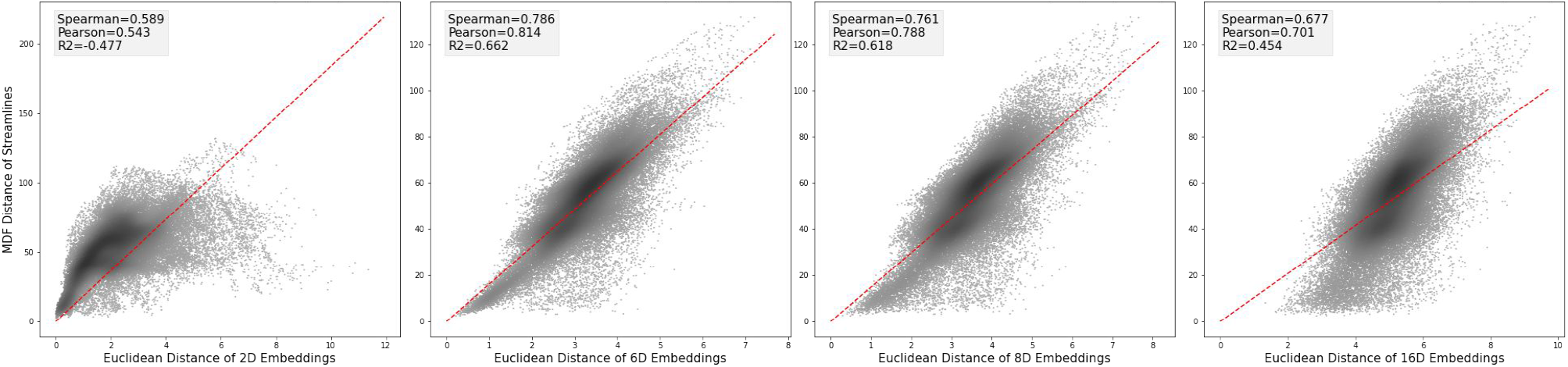
Distance correlation with latent space dimensions of size 2, 6, 8 and 16 calculated over 300 subsampled streamlines from the training set, using methods described in Detlefsen *et al*.^8^ The x-axis indicates Euclidean distances between pairs of embedded streamlines and the y-axis indicates MDF distances between the same pairs of streamlines in streamline space. Line fit with zero intercept is plotted as the red dashed line. As shown later (Figure 3), the embedding dimension 6 (2nd panel) preserves distances optimally. The Spearman *r*, Pearson *r* and *R*^2^ are marked in the legend.

**Figure 3.**
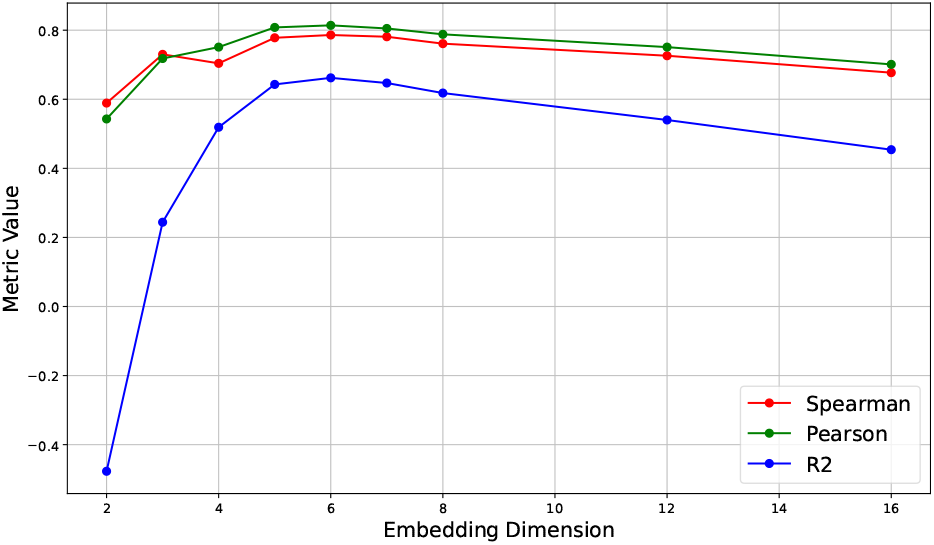
Plot of Pearson *r*, Spearman *r* and *R^2^* of line fit to evaluate metric preservation, in different models trained with 9 values of *N_z_*. The embedding of dimension 6 preserves distances optimally; Zhang *et al*.^30^ provides a theoretical analysis of optimal dimensions for discriminative classification with linear embeddings, in terms of the spectrum of the projection matrix, but in the ConvVAE case, this will depend on the number of bundles and empirical aspects of how they cluster.

### 3.2 Bundle Distance Preservation

Using ConvVAE with *N_z_* = 6, we further conduct distance correlation analyses for both bundles and their centroids to understand how bundle structural information is preserved in the latent space, as described in Section 2.3. The pairwise inter-bundle distance matrices are shown in Figure 4 and 5 respectively. The Mantel test (which evaluates the correlation between two pairwise distance matrices) was applied to distance matrices between bundles in the streamline and embedding spaces. For both bundles and centroids, this test was statistically significant (*p* = 0.01), with strong correlation *r_m_* = 0.84 and 0.98, as shown in Table 1. These results indicate that, in addition to the strong correlation from sampled streamlines, inter-bundle distances are well preserved in the ConvVAE latent space at the optimal embedding dimension of *N_z_* = 6.

**Figure 4.**
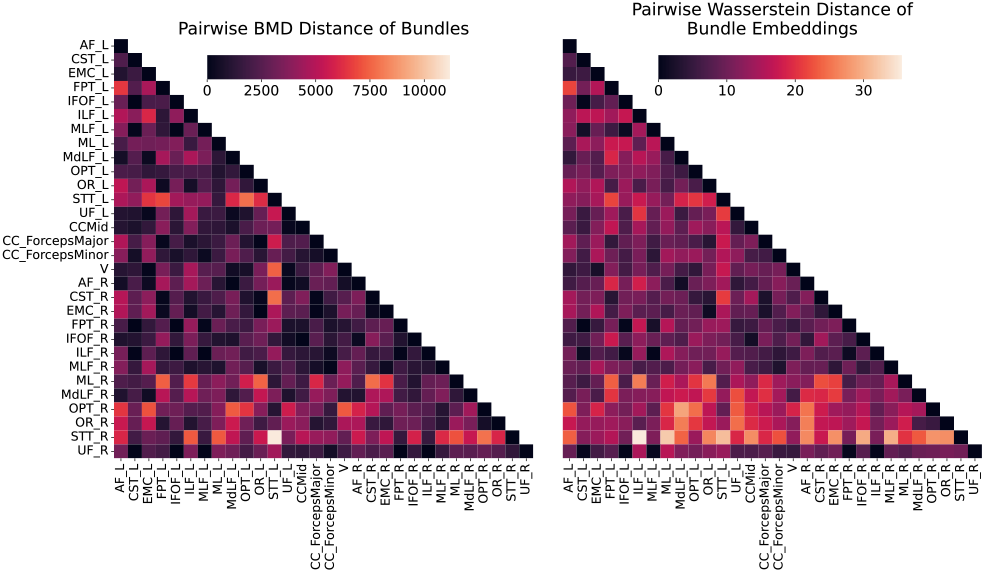
Pairwise inter-bundle distance matrix of bundles. BMD distances are calculated between (*X*(*b*)) and Wasserstein distances are calculated between (*Z*(*b*)) for *b* = 1, 2,…30.

**Figure 5.**
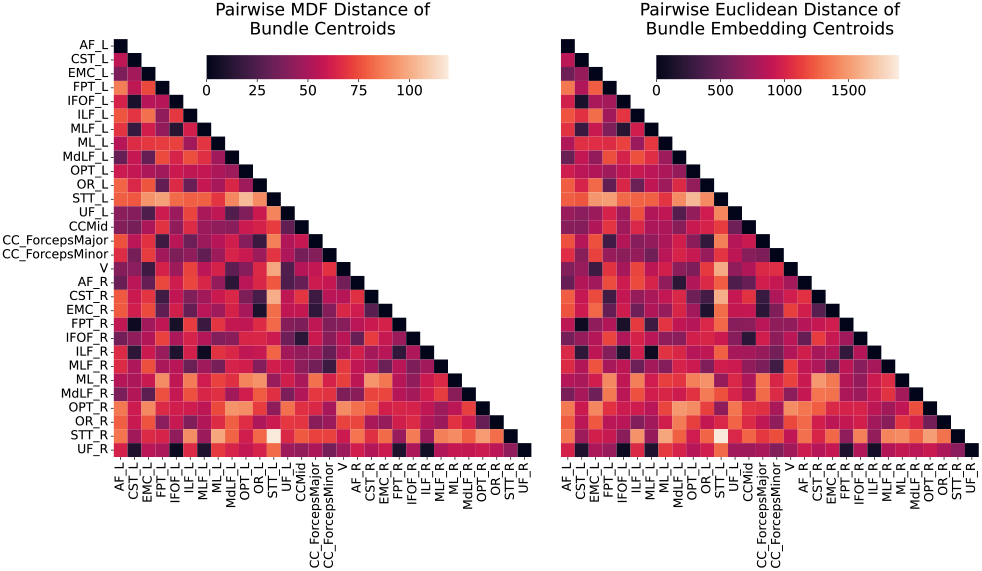
Pairwise inter-bundle distance matrix of bundle centroids. MDF distances are calculated between Centroid(*X*(*b*)) and Euclidean distances are calculated between Centroid(*Z*(*b*)) for *b* = 1, 2,…30.

### 3.3 Anomaly Detection

To detect anomalies in white matter bundles of participants with MCI and AD, we first conducted a group-wise comparison using weighted average MAE scores calculated from bundle reconstruction after controlling for age and sex. A bar plot of the difference between MAE scores per bundle per group for MCI and AD subjects and those from CN subjects are shown in Figure 6, where the bundles with significant results from the two-tailed independent sample t-tests are marked with an asterisk (*). In AD subjects, we found significant results at *α* = 0.5 after FDR correction in 6 bundles - the right middle longitudinal fasciculus (MdLF_R, *p* = 4.20×10^−5^), corpus callosum major (CC_ForcepsMajor, *p* = 4.85 × 10^-4^), left extreme capsule (EMC_L, *p* = 8.74 × 10^-4^), left arcuate fasciculus (AF_L, *p* = 4.20 × 10^-3^), right optic radiation (OR_R, *p* = 9.2 × 10^-3^), corpus callosum minor (CC_ForcepsMinor, *p* = 0.01) and corpus callosum middle sector (CCMid, *p* = 0.04). None of the bundles in MCI subjects showed detectable differences relative to those of CN subjects.

**Figure 6.**
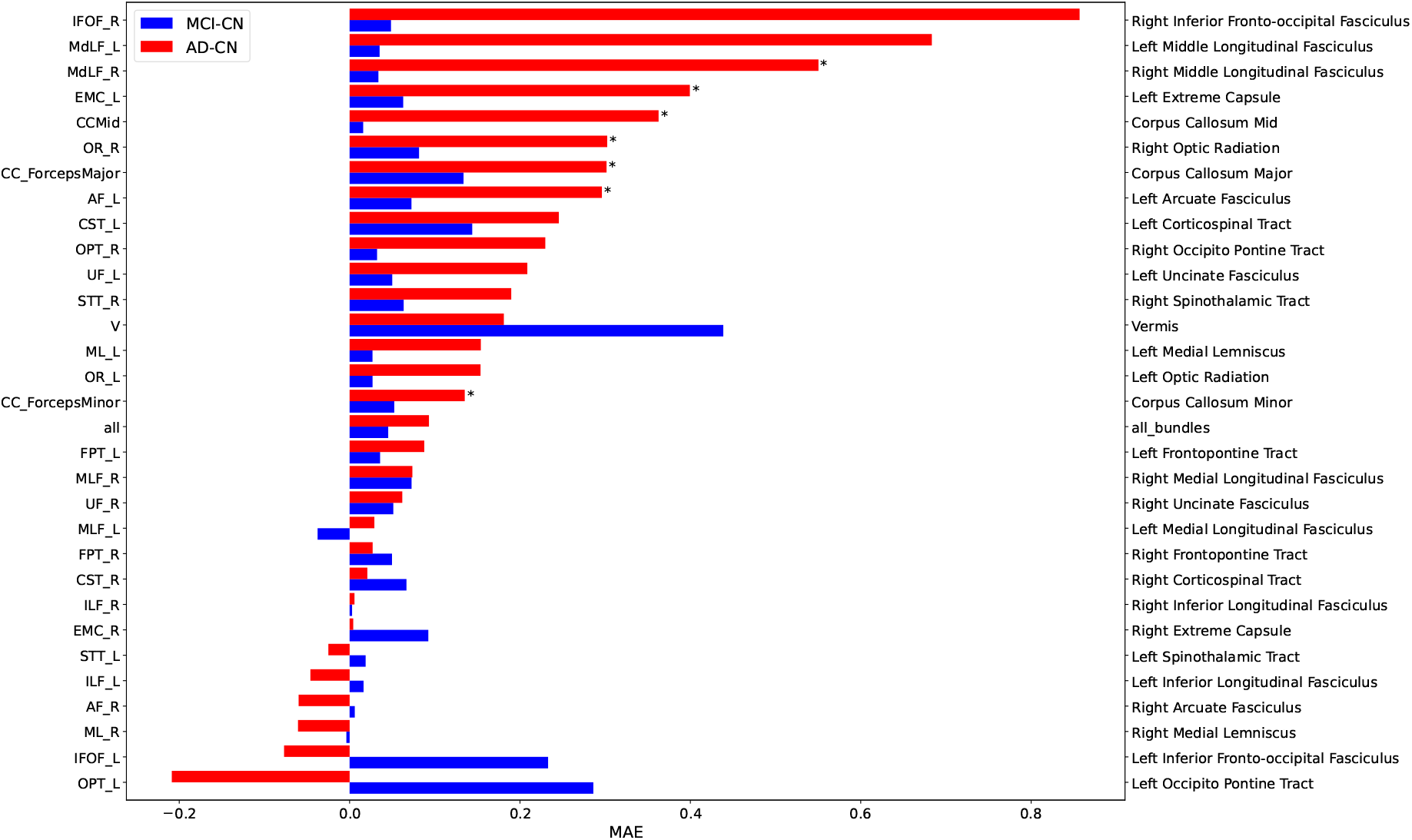
Bar plot of the difference of age and sex independent bundle MAE scores between MCI and AD subjects and those from CN subjects. Significant result from the two-tailed independent sample t-tests after FDR correction are marked with an asterisk (*). Full names of the bundles are indicated on the right y-axis.

To more closely evaluate the anomaly profile along each bundle in AD subjects versus CN subjects, we performed two-tailed independent sample t-tests with FDR correction on age and sex independent MAE scores for each of the 100 segments along the 7 bundles that were significant in the above group-wise comparison. The MdLF_R, CC_ForcepsMajor, AF_L, OR_R, CC_ForcepsMinor and CCMid tracts have at least one segment with statistically significant difference. Their MAE scores with 95% confidence interval and FDR-corrected — log_10_(*p*) values are plotted along with the tract colored by significance in Figures 7 and 8. The OR_R, MdLF_R and CC_ForcepsMajor tracts show high variation in the AD group compared to the CN and MCI groups. Notably, all three corpus callosum tracts show significant differences between AD and CN groups along the tracts, perhaps reflecting curvature differences due to ventricular dilation in dementia.^34^ The EMC_L tract, while having significant group differences overall, shows no significant difference along-tract due to age and sex effect. In the MdLF_R tract, segments 50-61 have the most pronounced group differences with FDR-corrected *p* = 0.02. The positions with significant group differences are aligned with those of high MAE scores in the MdLF_R tract, whereas in other tracts, not all points with significant difference have high MAE scores. While endpoints of tracts tend to have higher MAE scores due to greater inter-subject variation, MAE scores calculated from the ConvVAE still allow us to conduct group-wise comparisons and detect tract positions with significant differences.

**Figure 7.**
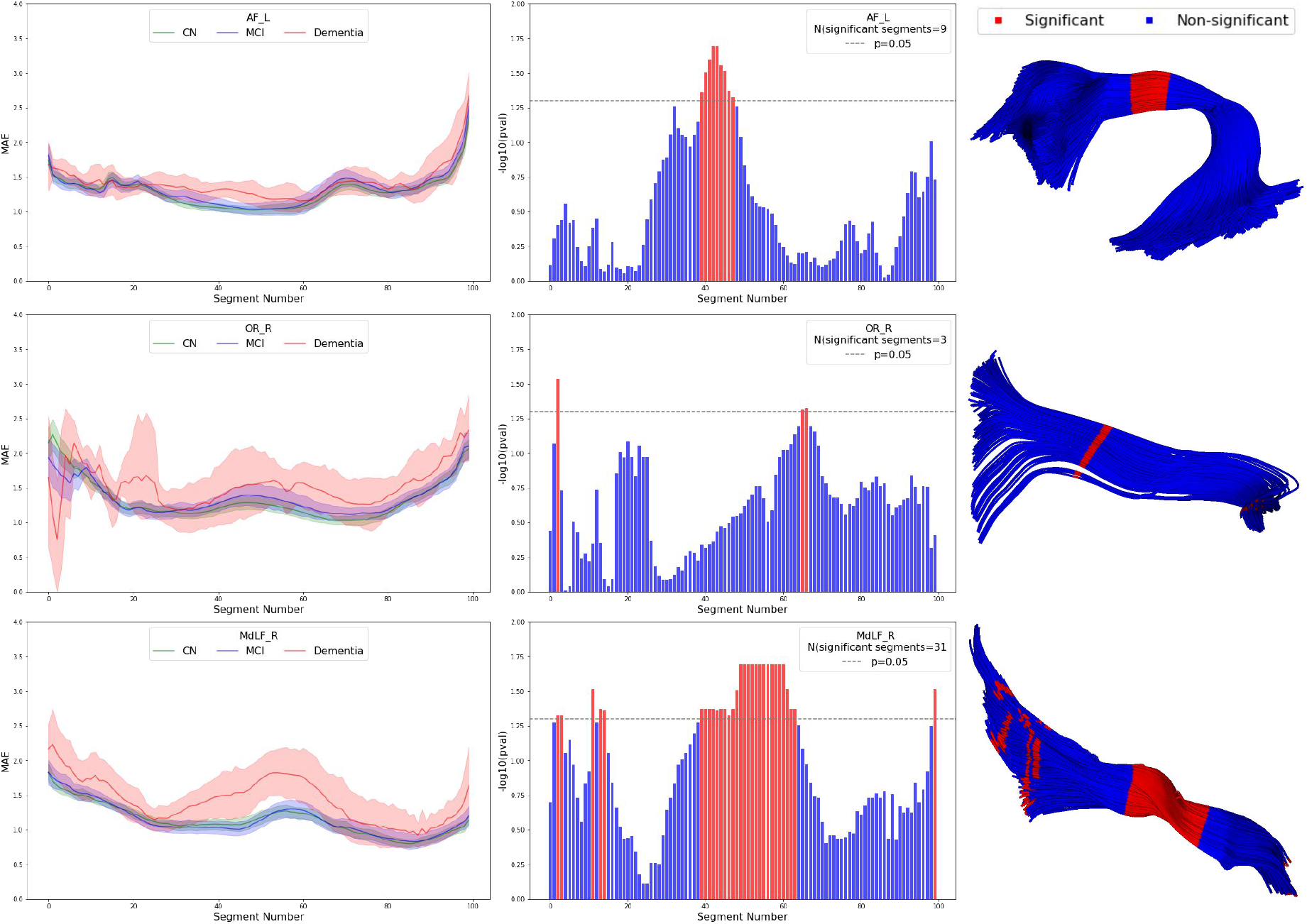
Segment-wise along-tract MAE scores after controlling for age and sex with 95% confidence intervals per diagnostic group for the AL_L, OR_R and MdLF_R tracts and FDR-corrected — log_10_(p) values plot from the 2-tailed independent sample t-test between control and AD are shown. The bundles are plotted on the atlas dataset colored by significance.

**Figure 8.**
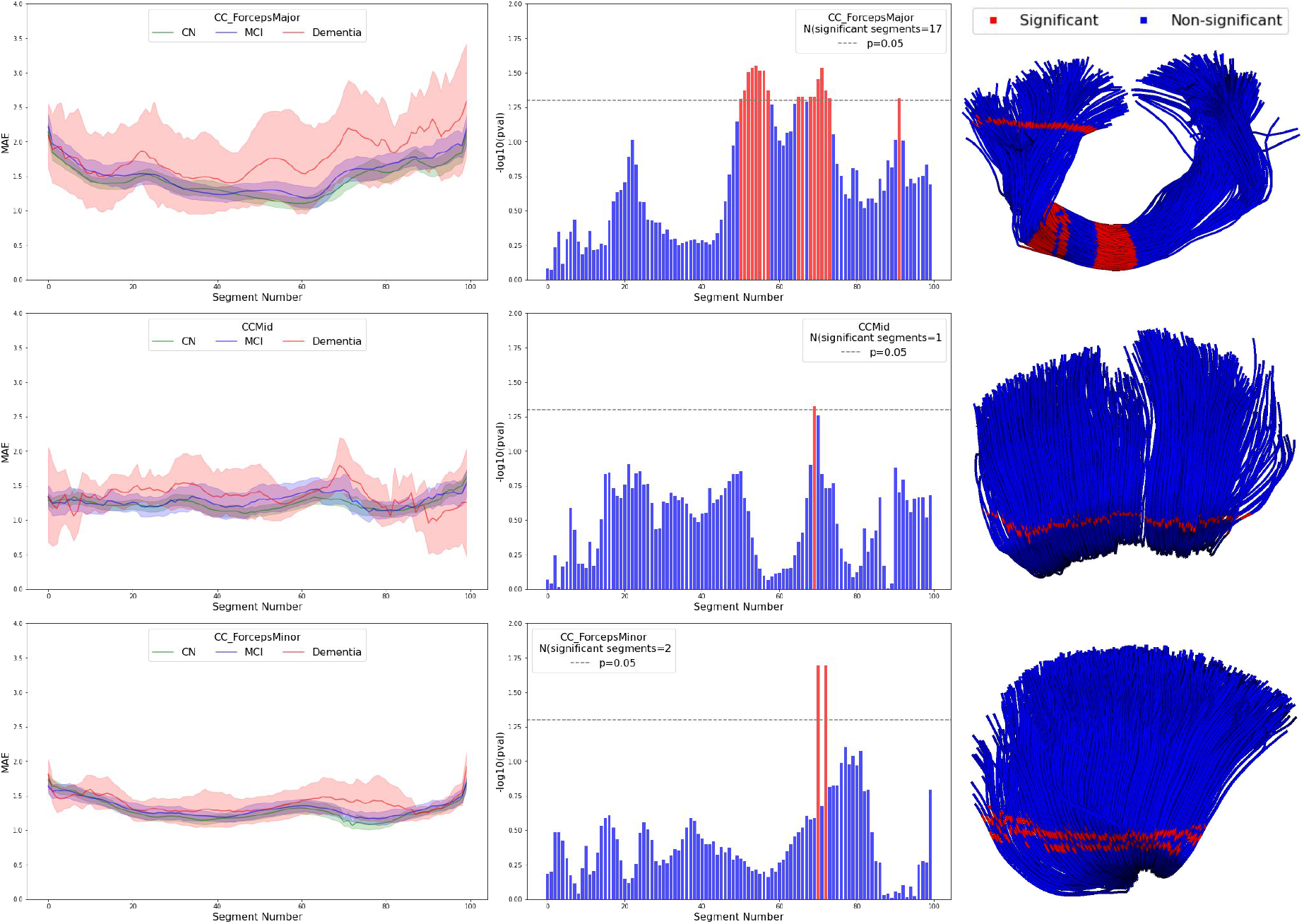
Segment-wise along-tract MAE scores after controlling for age and sex with 95% confidence intervals per diagnostic group for the CC_ForcepsMajor, CCMid and CC_ForcepsMinor tracts and FDR-corrected −log_10_ (*p*) values plot from the 2-tailed independent sample t-test between control and AD are shown. The bundles are plotted on the atlas dataset colored by significance.

## 4. DISCUSSION

Unsupervised representation learning methods have shown promise in learning embeddings from large datasets that enable downstream analysis, and lend themselves naturally to whole-brain tractography datasets with up to a million streamlines per subject. Applications include anomaly detection in individuals or groups, denoising, and quality control, as well as producing a more compact representation of the data for clustering and labeling. In this work using ConvVAE to encode bundle streamlines, we found that higher latent space dimensions lead to poorer distance preservation, potentially due to overfitting, while latent spaces of lower than 6 dimensions discard too much of the information needed to reconstruct tracts and their relative distances. Since our input data consists of bundle streamlines, we also designed inter-bundle distance evaluations to test whether global distances are preserved, using modality specific distance metrics.

We utilized our ConvVAE model to detect structural anomalies in white matter tracts of MCI and AD subjects. In the current formulation, structural anomalies are measured by the discrepancy between brains of people diagnosed with AD and MCI and normal brains using MAE scores computed over segments along the length of the tract. ConvVAE performs well for bundle reconstruction, preserving their shapes, orientations and locations in the brain, so we expect structural anomalies to be detected by MAE which uses reconstructed streamlines in its calculation. In addition to group analysis of bundles, the ConvVAE reconstructs streamlines, allowing us to compute along-tract measures. This approach help tease out significant group differences in points with high inter-subject variations inherent to many tractography methods.^35^

One limitation of our method is that ConvVAE with 1D convolutional layers can only take in equal-length inputs. Since not all streamlines have equal length, shorter streamlines are represented with more points, leading to bias against long streamlines which can affect downstream anomaly detection tasks. We plan in future work to adjust for streamline length and sampling to further improve reconstruction while preserving the quality of the embeddings. A second limitation is that our current work only flags geometric distortions along tracts and could be extended to map group differences in microstructural parameters, such as fractional anisotropy (FA) and mean diffusivity (MD) measures, which may be more sensitive to groups differences in MCI and AD.^13^ Our framework could be extended in several ways. First, we plan to train the model on a larger cohort with additional quality control on bundles, such as via the FiberNeat method.^9^ Evidently, the spikes in along-tract MAE in the OR_R tract (see Figure 7) are potentially due to outlier streamlines. Second, the current VAE embedding model uses a standard multidimensional Gaussian to determine the log-likelihood of the training data. *Contrastive learning* approaches, such as SimCLR^36^ and nearest-neighbor-based out-of-distribution based method,^37^ could instead be used to encourage mappings that cluster specific fiber types together in the latent space. In supervised embedding, labels (or numeric values) are leveraged so that similar points are closer together than they otherwise would be, and contrastive learning or semantic embedding could be used to pull streamlines from the same bundle together in the embedding space. This could allow direct multisubject registration and labeling of the embeddings for population analyses of microstructural and geometric parameters. Finally, a single VAE model for all tracts, used here, could be extended to a Gaussian mixture VAE^38^ to better capture the hierarchical structure of the bundles.

## 5. CONCLUSION

We propose a robust framework using Convolutional Variational Autoencoder (ConvVAE) to learn low-dimensional embeddings from data-intensive tractography data. We investigate the effect of latent space dimension on the quality of embeddings, and found that streamline distances as well as inter-bundle distances are strongly correlated with embedding distances at *N_z_* = 6. The generative model allows for inference on new data, and the smooth ConvVAE latent space enables meaningful decodings that can be used for downstream tasks. We trained our ConvVAE on data from healthy control subjects to detect structural anomalies in white matter tracts in patients with Alzheimer’s disease. The flexibility of ConvVAE facilitates group analysis of bundle difference. We identified 6 tracts with statistically significant group differences and specific locations along the length of the tracts with anomalies after controlling for age and sex effect despite large inter-subject variations. Given the increasing scale of neuroimaging studies and numerous tractography methods, our framework offers a robust, unsupervised method to study structural features of white matter tracts and conduct population analyses.

## REFERENCES

[1] Prasad, G., Nir, T. M., Toga, A. W., and Thompson, P. M., “Tractography Density and Network Measures in Alzheimer’s Disease,” Proceedings. IEEE Int. Symp. on Biomedical Imaging 2013, 692–695 (Apr. 2013).

[2] Amoroso, N., “Diffusion-weighted imaging (DWI) tractography and Alzheimer’s disease,” in [Diagnosis and Management in Dementia], 313–325, Elsevier (2020).

[3] Chandio, B. Q., Risacher, S. L., Pestilli, F., Bullock, D., Yeh, F.-C., Koudoro, S., Rokem, A., Harezlak, J., and Garyfallidis, E., “Bundle analytics, a computational framework for investigating the shapes and profiles of brain pathways across populations,” Scientific Reports 10, 17149 (Dec. 2020).

[4] Legarreta, J. H., Petit, L., Rheault, F., Theaud, G., Lemaire, C., Descoteaux, M., and Jodoin, P.-M., “Filtering in tractography using autoencoders (FINTA),” Medical Image Analysis 72, 102126 (Aug. 2021).

[5] Gupta, V., Thomopoulos, S. I., Corbin, C. K., Rashid, F., and Thompson, P. M., “Fibernet 2.0: An Automatic Neural Network Based Tool for Clustering White Matter Fibers in the Brain,” preprint, Neuroscience (Oct. 2017).

[6] Garyfallidis, E., Côté, M.-A., Rheault, F., Sidhu, J., Hau, J., Petit, L., Fortin, D., Cunanne, S., and Descoteaux, M., “Recognition of white matter bundles using local and global streamline-based registration and clustering,” NeuroImage 170, 283–295 (Apr. 2018).

[7] Manjón, J. V., Coupé, P., Concha, L., Buades, A., Collins, D. L., and Robles, M., “Diffusion Weighted Image Denoising Using Overcomplete Local PCA,” PLoS ONE 8, e73021 (Sept. 2013).

[8] Detlefsen, N. S., Hauberg, S., and Boomsma, W., “Learning meaningful representations of protein sequences,” Nature Communications 13, 1914 (Dec. 2022).

[9] Chandio, B. Q., Chattopadhyay, T., Owens-Walton, C., Villalon Reina, J. E., Nabulsi, L., Thomopoulos, S. I., Garyfallidis, E., and Thompson, P. M., “FiberNeat: unsupervised streamline clustering and white matter tract filtering in latent space,” preprint, Neuroscience (Oct. 2021).

[10] Zhong, S., Chen, Z., and Egan, G., “Auto-encoded Latent Representations of White Matter Streamlines for Quantitative Distance Analysis,” Neuroinformatics (June 2022).

[11] Steck, H., “Autoencoders that don’ t overfit towards the Identity,” in [Advances in Neural Information Processing Systems], Larochelle, H., Ranzato, M., Hadsell, R., Balcan, M. F., and Lin, H., eds., 33, 19598–19608, Curran Associates, Inc. (2020).

[12] Feng, Y., Chandio, B. Q., Chattopadhyay, T., Thomopoulos, S. I., Owens-Walton, C., Jahanshad, N., Gary-fallidis, E., and Thompson, P. M., “Deep generative model for learning tractography streamline embeddings based on convolutional variational autoencoder,” in [International Society for Magnetic Resonance Imaging (ISMRM)], (2022).

[13] Zavaliangos-Petropulu, A., Nir, T. M., Thomopoulos, S. I., Reid, R. I., Bernstein, M. A., Borowski, B., Jack Jr., C. R., Weiner, M. W., Jahanshad, N., and Thompson, P. M., “Diffusion MRI Indices and Their Relation to Cognitive Impairment in Brain Aging: The Updated Multi-protocol Approach in ADNI3,” Frontiers in Neuroinformatics 13, 2 (Feb. 2019).

[14] Garyfallidis, E., Brett, M., Amirbekian, B., Rokem, A., van der Walt, S., Descoteaux, M., Nimmo-Smith, I., and Dipy Contributors, “Dipy, a library for the analysis of diffusion MRI data,” Frontiers in Neuroinformatics 8 (Feb. 2014).

[15] Veraart, J., Fieremans, E., and Novikov, D. S., “Diffusion MRI noise mapping using random matrix theory: Diffusion MRI Noise Mapping,” Magnetic Resonance in Medicine 76, 1582–1593 (Nov. 2016).

[16] Veraart, J., Novikov, D. S., Christiaens, D., Ades-aron, B., Sijbers, J., and Fieremans, E., “Denoising of diffusion MRI using random matrix theory,” NeuroImage 142, 394–406 (Nov. 2016).

[17] Kellner, E., Dhital, B., Kiselev, V. G., and Reisert, M., “Gibbs-ringing artifact removal based on local subvoxel-shifts: Gibbs-Ringing Artifact Removal,” Magn. Res. Medicine 76, 1574–1581 (Nov. 2016).

[18] Andersson, J. L. and Sotiropoulos, S. N., “An integrated approach to correction for off-resonance effects and subject movement in diffusion MR imaging,” NeuroImage 125, 1063–1078 (Jan. 2016).

[19] Andersson, J. L. R., Graham, M. S., Zsoldos, E., and Sotiropoulos, S. N., “Incorporating outlier detection and replacement into a non-parametric framework for movement and distortion correction of diffusion MR images,” NeuroImage 141, 556–572 (Nov. 2016).

[20] Andersson, J. L., Graham, M. S., Drobnjak, I., Zhang, H., Filippini, N., and Bastiani, M., “Towards a comprehensive framework for movement and distortion correction of diffusion MR images: Within volume movement,” NeuroImage 152, 450–466 (May 2017).

[21] Tournier, J.-D., Calamante, F., and Connelly, A., “Robust determination of the fibre orientation distribution in diffusion MRI: non-negativity constrained super-resolved spherical deconvolution,” NeuroImage 35, 1459–1472 (May 2007).

[22] Girard, G., Whittingstall, K., Deriche, R., and Descoteaux, M., “Towards quantitative connectivity analysis: reducing tractography biases,” NeuroImage 98, 266–278 (Sept. 2014).

[23] Kingma, D. P. and Welling, M., “Auto-Encoding Variational Bayes,” (May 2014). arXiv: 1312.6114 [cs, stat].

[24] Kingma, D. P. and Ba, J., “Adam: A Method for Stochastic Optimization,” (Jan. 2017). arXiv:1412.6980 [cs].

[25] Mikolov, T., “Statistical language models based on neural networks,” (2012).

[26] Garyfallidis, E., Brett, M., Correia, M. M., Williams, G. B., and Nimmo-Smith, I., “QuickBundles, a Method for Tractography Simplification,” Frontiers in Neuroscience 6, 175 (2012).

[27] Garyfallidis, E., Wassermann, D., and Descoteaux, M., “Direct native-space fiber bundle alignment for group comparisons,” in [International Society for Magnetic Resonance Imaging (ISMRM)], (2014).

[28] Kantorovich, L. V., “Mathematical Methods of Organizing and Planning Production,” Management Science 6, 366–422 (July 1960).

[29] Mantel, N., “The detection of disease clustering and a generalized regression approach,” Cancer Research 27, 209–220 (Feb. 1967).

[30] Zhang, W., Xue, X., Sun, Z., Guo, Y.-F., and Lu, H., “Optimal dimensionality of metric space for classification,” in [Proceedings of the 24th international conference on Machine learning - ICML ’07], 1135–1142, ACM Press, Corvalis, Oregon (2007).

[31] Baur, C., Denner, S., Wiestler, B., Albarqouni, S., and Navab, N., “Autoencoders for Unsupervised Anomaly Segmentation in Brain MR Images: A Comparative Study,” (Apr. 2020). arXiv:2004.03271 [cs, eess].

[32] Siddalingappa, R. and Kanagaraj, S., “Anomaly Detection on Medical Images using Autoencoder and Convolutional Neural Network,” Int. J. Advanced Computer Science and Applications 12(7) (2021).

[33] Chamberland, M., Genc, S., Tax, C. M. W., Shastin, D., Koller, K., Raven, E. P., Cunningham, A., Doherty, J., van den Bree, M. B. M., Parker, G. D., Hamandi, K., Gray, W. P., and Jones, D. K., “Detecting microstructural deviations in individuals with deep diffusion MRI tractometry,” Nature Computational Science 1, 598–606 (Sept. 2021).

[34] Ardekani, B. A., Bachman, A. H., Figarsky, K., and Sidtis, J. J., “Corpus callosum shape changes in early Alzheimer’s disease: an MRI study using the OASIS brain database,” Brain Structure & Function 219, 343–352 (Jan. 2014).

[35] Rheault, F., Poulin, P., Valcourt Caron, A., St-Onge, E., and Descoteaux, M., “Common misconceptions, hidden biases and modern challenges of dMRI tractography,” J. Neural Engineering 17, 011001 (Feb. 2020).

[36] Chen, T., Kornblith, S., Norouzi, M., and Hinton, G., “A Simple Framework for Contrastive Learning of Visual Representations,” arXiv:2002.05709 [cs, stat] (June 2020). arXiv: 2002.05709.

[37] Sun, Y., Ming, Y., Zhu, X., and Li, Y., “Out-of-Distribution Detection with Deep Nearest Neighbors,” (June 2022). arXiv:2204.06507 [cs].

[38] Jiang, Z., Zheng, Y., Tan, H., Tang, B., and Zhou, H., “Variational Deep Embedding: An Unsupervised and Generative Approach to Clustering,” in [Proceedings of the Twenty-Sixth International Joint Conference on Artificial Intelligence], 1965–1972, International Joint Conferences on Artificial Intelligence Organization, Melbourne, Australia (Aug. 2017).

